# The lipidome of posttraumatic stress disorder

**DOI:** 10.1101/2024.02.23.581833

**Authors:** Aditi Bhargava, Johannes D. Knapp, Oliver Fiehn, Thomas C. Neylan, Sabra S. Inslicht

## Abstract

Posttraumatic stress disorder (PTSD) can develop after trauma exposure. Some studies report that women develop PTSD at twice the rate of men, despite greater trauma exposure in men. Lipids and their metabolites (lipidome) regulate a myriad of key biological processes and pathways such as membrane integrity, oxidative stress, and neuroinflammation in the brain by maintaining neuronal connectivity and homeostasis. In this study, we analyzed the lipidome of 40 individuals with PTSD and 40 trauma-exposed non-PTSD individuals. Plasma samples were analyzed for lipidomics using Quadrupole Time-of-Flight (QToF) mass spectrometry. Additionally, ∼ 90 measures were collected, on sleep, mental and physical health indices. Sleep quality worsened as PTSD severity increased in both sexes. The lipidomics analysis identified a total of 348 quantifiable known lipid metabolites and 1951 lipid metabolites that are yet unknown; known metabolites were part of 13 classes of lipids. After adjusting for sleep quality, in women with PTSD, only one lipid subclass, phosphatidylethanolamine (PE) was altered, whereas, in men with PTSD, 9 out of 13 subclasses were altered compared to non-PTSD women and men, respectively. Severe PTSD was associated with 22% and 5% of altered lipid metabolites in men and women, respectively. Of the changed metabolites, only 0.5% measures (2 PEs and cholesterol) were common between women and men with PTSD. Several sphingomyelins, PEs, ceramides, and triglycerides were increased in men with severe PTSD. The triglycerides and ceramide metabolites that were most highly increased were correlated with cholesterol metabolites and systolic blood pressure in men but not always in women with PTSD. Alterations in triglycerides and ceramides are linked with cardiac health and metabolic function in humans. Thus, disturbed sleep and higher weight may have contributed to changes in the lipidome found in PTSD.

## Introduction

Posttraumatic stress disorder (PTSD) is a psychiatric condition that may develop after exposure to an actual or threatened death, serious injury, or sexual violence. PTSD may be characterized by: (i) reexperiencing (e.g., intrusive thoughts, nightmares, flashbacks); (ii) avoidance; (iii) negative changes in cognition and mood (hopelessness, lack of emotions), and (iv) hyperarousal (trouble sleeping, risky or destructive behavior, angry outbursts) (DSM-5)^1^. Disturbed sleep is one of the most common complaints among individuals with PTSD^2^. Lower slow wave sleep duration and delta-band spectral power are more pronounced in men than women^3^, and correlate with PTSD status. In contrast, greater rapid eye movement sleep is found in women with PTSD compared to healthy controls, a difference not seen in men^3^. PTSD is also known to affect physical health and has been associated with greater inflammation, metabolic syndrome, gastrointestinal illness, and even early mortality^4–13^. It is possible that disturbed sleep may play a role in these health impacts since sleep duration correlates with metabolic risk in PTSD^14^.

Epidemiological evidence suggests that women develop PTSD at twice the rate of men, despite greater trauma exposure in men^15,16^. Women are also at increased risk for stress-related physical comorbidities, including inflammatory, metabolic, and GI disorders^15,17–19^. Although some have suggested that greater exposure to interpersonal violence may contribute to higher rates of PTSD in women; evidence also implicates sex differences in the molecular mechanisms involved in stress regulation and disease processes. Trauma exposure can have significant effects on molecular, biochemical, and cellular systems that are associated with a complex array of PTSD symptoms and physiological comorbidities^20,21^. Lipids are emerging as an important contributor of health of the brain but the relationship between PTSD severity and lipid metabolites is unknown. While prior studies have found alterations in neuroendocrine, immune, and aging processes in PTSD^4–13^, our understanding of the role of metabolite disturbances in PTSD is limited.

The structure and function of the complete set of lipids in each cell or organism is referred to as the “Lipidome”. Several classes of lipids that include fatty acids, diacylglycerols, triglycerides, phospholipids, sphingomyelin, ceramides, and acylcarnitine comprise the lipidome (Fig. 1a)^22^. Most of the classes of lipids are derived from fatty acids with Acyl-CoA as a key intermediary (Fig. 1). Phospholipids form the structural basis of all cellular membranes and account for nearly 25% of the dry weight of an adult human brain. Phosphatidylethanolamine (PE) together with phosphatidylcholine (PC), phosphatidylserine (PS) and phosphatidylinositol (PI) form the backbone of most biological membranes. The relative proportions of lipid subclasses are maintained at a steady state under homeostasis. Phospholipid subclasses regulate critical physiological activities such as cell signaling, membrane structure, fluidity, permeability, organelle, and immune functions. Membrane fluidity, especially for neuronal cells is key for their structure and function and is determined by the presence phospholipid subclasses and their topological distribution within the cell and organelle membranes^23^. The most abundant phospholipid subclass in cell membranes is phosphatidylcholine. PCs serve two key functions-determine membrane fluidity and storage of neurotransmitter choline. Membranes with PC as predominant composition are devoid of any curvature due to unique molecular geometry, and are typically fluid at room temperature^24^. Since the ratio of PC to other phospholipids determine membrane shape and permeability, altered ratio can lead to neuronal, cellular and organelle signaling dysfunction. PC also serves as an essential reservoir for storing choline, a precursor for the neurotransmitter acetylcholine and is essential for proper brain/neuronal function^25^. While altered levels of PC are seen in individuals with traumatic brain injury and is associated with impaired cholinergic neurotransmission and impaired neurogenesis^26^, it is unclear if the same class of lipids are also altered in individuals with PTSD.

**Figure 1.**
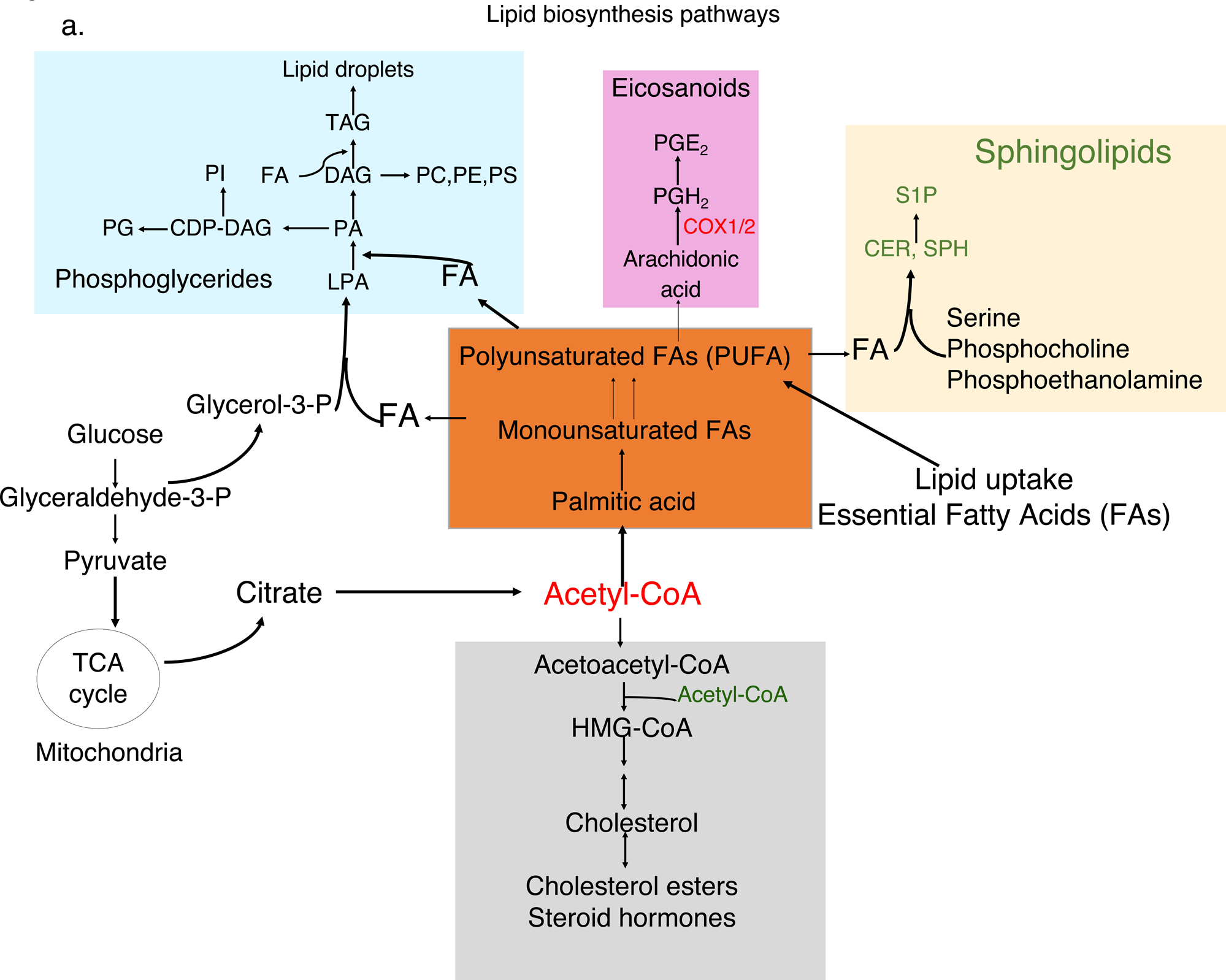
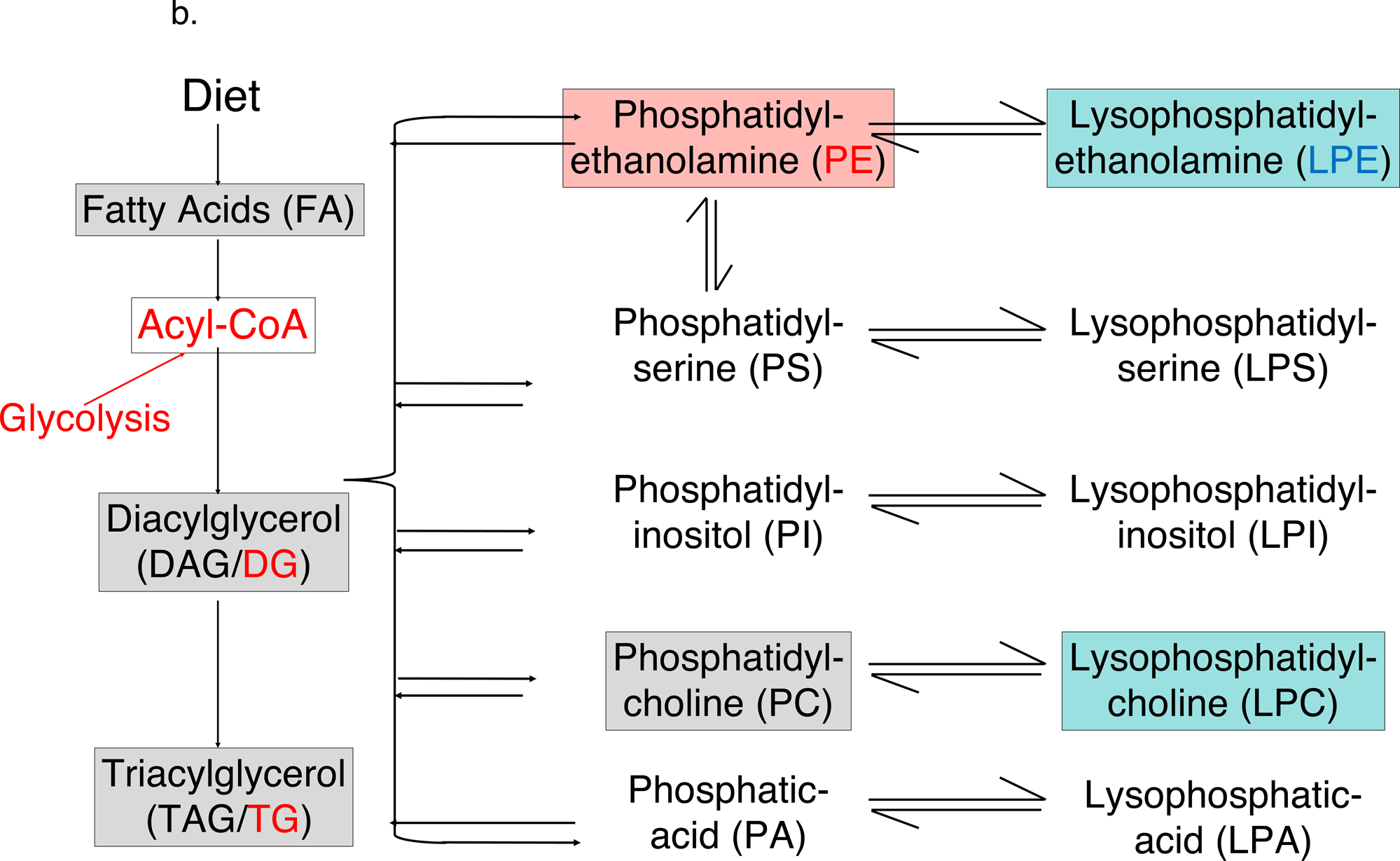
Lipid biosynthesis pathways and lipid metabolites. (**a**). Schematic of pathways involved in the synthesis of various lipid subclasses from Acetyl-Co-A. Glucose is converted to Acetyl-Co-A via glycolysis in the mitochondrial membrane (tricarboxylic acid cycle: TCA). Introduction of a double bond in the Δ 9 position of the acyl chain and subsequent elongation of the carbon chain leads to the generation of mono and poly unsaturated fatty acids (PUFA), respectively. Essential omega FA cannot be synthesized by human cells and must be obtained from the diet. Sphingolipids contain polar heads derived from serine, phosphocholine, or phosphoethanolamine. Cholesterol is also synthesized from Acetyl-Co-A and is the structural backbone and starting material for all steroid hormone biosynthesis. (**b**). Biosynthesis pathway of 13 major lipid subclasses identified using MToF mass spectrometry. COX1/2: prostaglandin-endoperoxide synthase. CDP-DAG: cytidine diphosphate-diacylglycerol; CER: ceramide; DAG/DG: diacylglycerol; FA: fatty acid; LPA: lysophosphatidic acid; LPC: lysophosphatidylcholine; LPE: lysophosphatidylethanolamine; LPS: lysophosphatidylserine; PA: phosphatidic acid; PC: phosphatidylcholine; PE: phosphatidylethanolamine; PG: phosphatidylglycerol; PGE_2_: prostaglandin E_2_; PGH_2_: prostaglandin H_2_; PI: phosphatidylinositol; PIPx: phosphatidylinositol phosphate; PS: phosphatidylserine; S1P: sphingosine-1-phosphate; SP:, sphingosine; TAG/TG: triacylglyceride. Adapted from Baenke et al.

Lysophosphatidylcholine (LPC) or lysolecithins are derived from PC (Fig. 1b) due to cleavage by enzyme phospholipase A_2_ and/or by the transfer of fatty acids to free cholesterol via lecithin-cholesterol acyltransferase. Increased levels of LPCs disrupt mitochondrial membrane integrity and dysregulate cytochrome C release in hepatocytes to modulate cholesterol biosynthesis. High LPCs levels are associated with several pathologies such as cardiovascular diseases and diabetes. LPC modulate several endothelial functions; LPCs activate endothelial cells to induce chemokine expression and release, impairs arterial relaxation, increases oxidative stress, and inhibits endothelial cell migration and proliferation [23,24]. LPCs may serve as a group of proinflammatory lipids that are involved in the pathogenesis of central nervous system-associated disorders^23^. Phospholipids also modulate adaptive immune system by altering function of both B and T lymphocytes. Cell membrane assembly and organelle biogenesis require phospholipids as raw materials in activated B lymphocytes^27^. Neuronal activity and immune function are two key physiological processes that are altered in people with PTSD. The subclasses of phospholipids that are altered in men and women with PTSD is largely not known. PE have an essential role in chaperoning membrane proteins to their folded state; PEs catalyze the conversion of prions from the nontoxic to the toxic conformation. PE initiate autophagosome formation by covalently attaching to the autophagy protein Atg8. PEs are associated with ER stress associated with diabetes and neurodegeneration. PE with an unsaturated acyl chain is known to facilitate ferroptosis.

Mass spectrometry has allowed for the discovery of novel pathways using an unbiased method to examine multiple analytes simultaneously. Metabolomics is a global and unbiased approach to understanding regulation of metabolic pathways and networks of physiologically relevant interactions. The metabolome is regulated by gene-environment interactions and reflects the intermediary state between genotype and phenotype. Gene mutations, single nucleotide polymorphisms, and mutations in proteins are associated with PTSD, but none of these alone explains the complex manifestation of PTSD and comorbid health conditions. A multi-omics approach has been used to identify potential biomarkers that range from DNA methylation, proteins, miRNA, lipids, and other metabolites in warzone male veterans with PTSD^28^. Metabolomic profiling has also led to identification of key differences in glycolysis and fatty acid pathways that were associated with mitochondrial dysfunction in men with PTSD^29^. Characterization of metabolomics can help elucidate new discovery of yet unknown biological mechanisms of disease.

We are not aware of any study that has systematically ascertained sex differences in the status of lipid metabolites in serum samples of women and men with PTSD and trauma-exposed non-PTSD individuals. In this study, we aimed to examine alterations of lipid metabolites and the contribution of sleep measures in both men and women with PTSD using integrated systems analysis approach.

## Results

### Demographic Data and Clinical Characteristics

By design, PTSD and control subjects were sex-and age-matched, nor were there significant differences in education, or race/ethnicity across all four groups^14^. In our cohort, men and women with PTSD did not differ in terms of CAPS scores, rates of current MDD, or history of childhood trauma (defined by the presence of two or more categories of childhood trauma as compared to one or none)^3^. Eleven control subjects reported a lifetime history of a traumatic criterion A1 event, but all had current CAPS scores of zero and none had a lifetime history of PTSD. However, women with PTSD had higher PCL-C (avoidance) scores than men with PTSD (Fig. 2a). Additionally, none of the control subjects reported a history of two or more categories of childhood trauma. There were no differences between PTSD and control women in use of hormonal birth control or group differences in smoking of tobacco. Men with moderate and severe PTSD symptoms had higher BMI scores compared with women with similar scores and men with low PCL score (controls) had the lowest BMI (Fig. 2a). Clinical data from a total of 98 different measures, including sleep measures (Supplementary Table S1), were analyzed together with lipid metabolites in an integrated systems approach.

**Figure 2.**
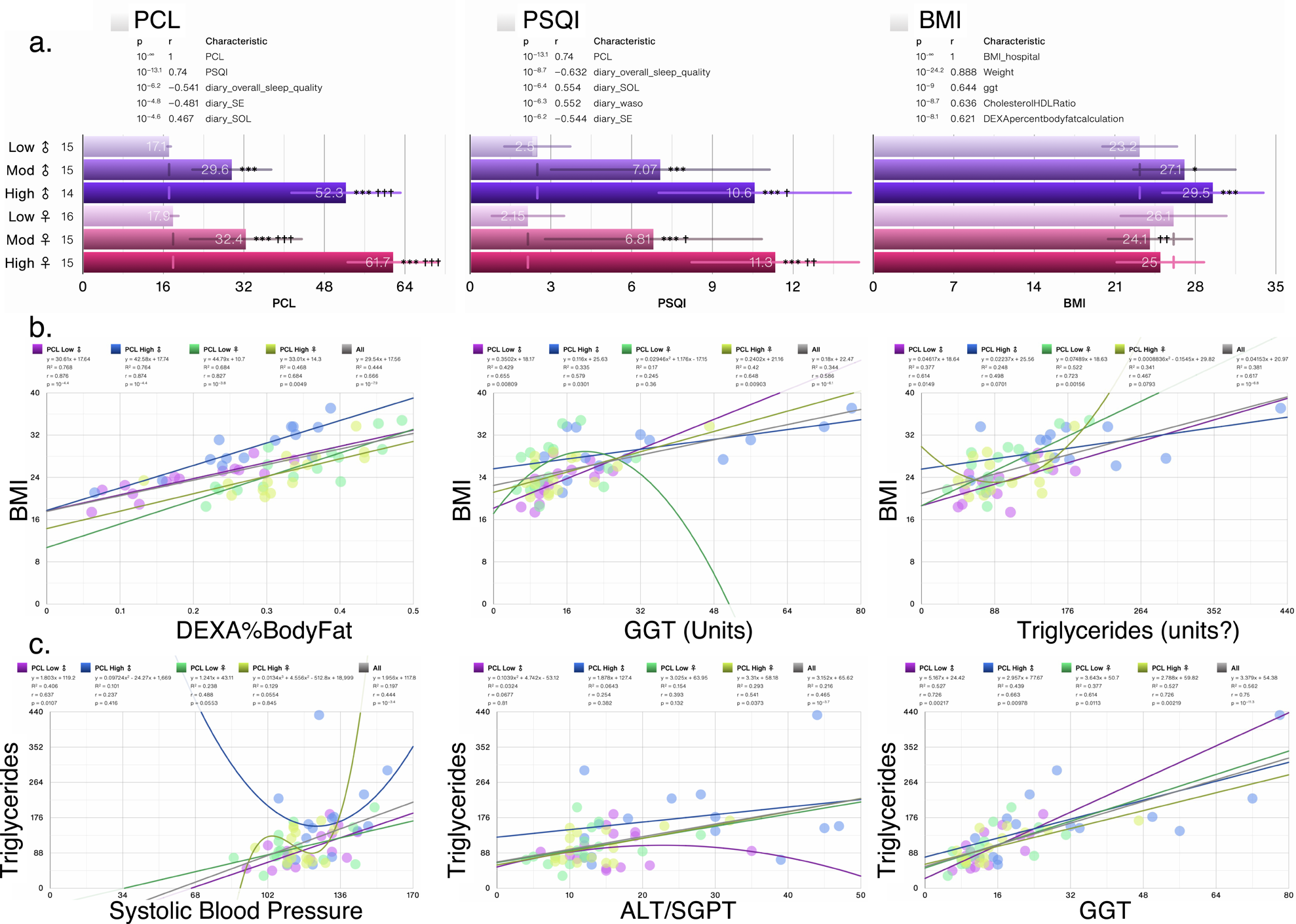
Clinical characteristics of PTSD cohort. (**a**) Bar charts showing the average BMI, PCL and PSQI scores in patient subgroups generated using 0-30% (Low PCL), 33-67% (Moderate PCL) and 67-100% (High PCL) scores. (**b-c**) Scatter plots for the four measures most significantly correlated with top 4 measures in the dataset (including clinical and lipid metabolites) Trendlines, or an n^th^-degree polynomial trendline if its goodness-of-fit is either 50% greater than, or if it explains at least half of the variance not explained by the (n – 1)^th^-degree polynomial, were fit to the data. R^2^, r and p denote goodness-of-fit, Pearson’s correlation coefficient and significance of the correlation, respectively. Numbers on X-axis in bar charts in (**a**) denote the actual number of patients in which the measures were detected. N/group: Low PCL ♀: 16; Moderate (Mod) PCL ♀: 15; High PCL♀: 15; Low PCL ♂: 15; Mod PCL ♂: 15; High PCL ♂: 14. GGT: glutamyltransferase; ALT/SGPT: serum glutamic-pyruvic transaminase

### Correlation of lipid subclasses with anthropomorphic and clinical measures of BMI and triglycerides

A PCL score range of 17-28 is considered as cut off as individuals within this score range show little-to-no PTSD symptoms, those with scores 28-29 show some PTSD symptoms, whereas scores >45 are considered as high severity of PTSD symptoms. In our cohort, average PCL score in women and men with no PTSD symptoms was 17.1; the average PCL score was 29.6 and 32-4 in men and women, respectively, with moderate PTSD symptoms, whereas individuals with high-to severe PTSD symptoms had a PCL score of >52.3 (Fig. 2a). Perceived sleep quality (PSQI) score was worst in men and women with PCL scores of >52 (Fig. 2a). BMI was highly correlated with % body fat as ascertained using DEXA scan (Fig. 2b). BMI was also highly correlated in men and women with high PCL score, but no correlation with BMI was evident in women with low PCL score (Fig. 2b). Correlation between BMI and clinical measure of triglyceride was also evident in men and women (Fig. 2b). While triglyceride levels were correlated with systolic blood pressure levels in men and women with low PCL score, this correlation was lost in all individuals as PTSD symptoms worsened (Fig. 2c). Interestingly, triglyceride levels correlated with plasma gamma-glutamyltransferase (GGT) levels, but serum glutamic-pyruvic transaminase (ALT/SGPT) levels were only correlated in women with high PCL scores. Both GGT and ALT functions are indicator of liver dysfunction, and GGT levels were associated with BMI, blood pressure, and triglycerides in the Framingham Heart Study^30,31^.

We next ascertained which lipid subclasses associated with BMI and weight. Of the 13 lipid subclasses, a total of 9 associated with BMI in men and only 3 in women, whereas only PI associated with BMI in men and 2 (fatty acids (FA) and triglycerides (TG)) in women (Table 1). In BMI unadjusted analysis, total blood cholesterol and calculated low-density lipoprotein (LDL) cholesterol exhibited significant correlation with 12/13 and 11/13 lipid subclasses, respectively in sex aggregated analysis (Table 2) with cholesterol esters and sphingomyelin subclasses exhibiting r>0.82 in both women and men (Table 1). Triglycerides measured with mass spectrometry in our dataset correlated highly with clinical measure of TGs measured by routine assays (r=>0.95) in both women and men (Table 2). TGs in our dataset correlated very strongly with calculated VLDL (r=>0.94) and cholesterol:HDL ratio (r=>0.61) in both sexes, but less strongly with total cholesterol, HDL and VLDL levels (Table 2). In addition to TG, 8 other lipid subclasses correlated with blood TG levels in men and 5 subclasses in women with DG (r=> 0.89) and ceramides exhibiting strong correlation. In women, except for LPC, all other subclasses associated with VLDL levels (Table 2).

**Table 1.**
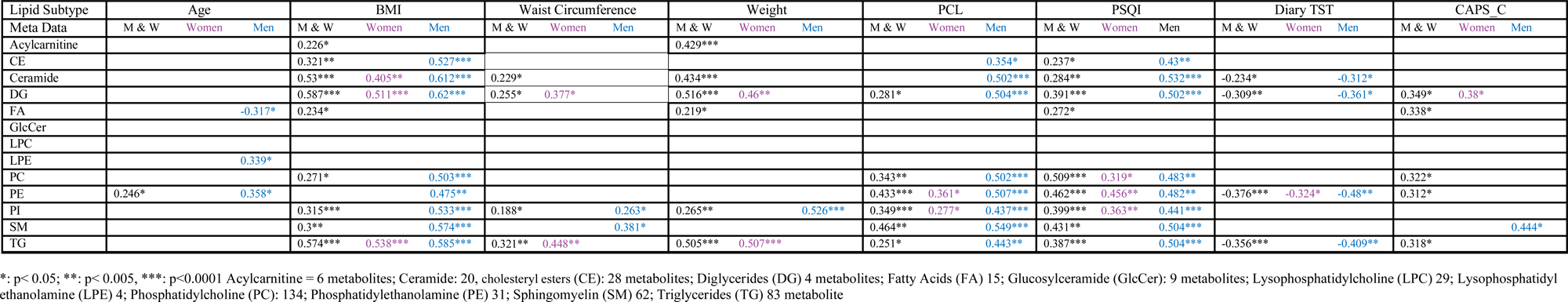
Lipid Metabolite subgroups r values-correlation with anthropometric and sleep measures.

**Table 2.** Lipid Metabolite subgroups r values-correlation with lab measures of Cholesterol, Triglycerides and Blood Pressure.

| Lipid Subtype | Cholesterol (Total) |  |  | Cholesterol HDL |  |  | Cholesterol LDL |  |  | Cholesterol VLDL |  |  | Cholesterol:HDL Ratio |  |  | Triglycerides (clinical) |  |  | Diastolic Blood Pressure |  |  | Systolic Blood Pressure |  |  |
| --- | --- | --- | --- | --- | --- | --- | --- | --- | --- | --- | --- | --- | --- | --- | --- | --- | --- | --- | --- | --- | --- | --- | --- | --- |
| Meta Data | M & W | Women | Men | M & W | Women | Men | M & W | Women | Men | M & W | Women | Men | M & W | Women | Men | M & W | Women | Men | M & W | Women | Men | M & W | Women | Men |
| Acylcarnitine | 0.384*** | 0.432** | 0.392* |  |  |  | 0.318** | 0.384* |  | 0.352** |  | 0.361* | 0.316** |  |  | 0.384*** |  | 0.4* |  |  |  |  |  |  |
| CE | 0.809*** | 0.8*** | 0.838*** | 0.26* | 0.403** |  | 0.732*** | 0.763*** | 0.743*** | 0.292** |  | 0.395* | 0.419*** | 0.323* | 0.592*** | 0.357** |  | 0.504*** | 0.408*** | 0.357* | 0.501*** | 0.256* |  | 0.333* |
| Ceramide | 0.72*** | 0.71*** | 0.737*** |  |  |  | 0.625*** | 0.689*** | 0.571*** | 0.659*** | 0.518*** | 0.783*** | 0.605*** | 0.49*** | 0.679*** | 0.719*** | 0.52*** | 0.842*** | 0.417*** | 0.342* | 0.458** | 0.386*** | 0.305* | 0.42** |
| DG | 0.505*** | 0.5*** | 0.523*** | -0.252* |  | -0.307* | 0.333** | 0.392* |  | 0.894*** | 0.891*** | 0.899*** | 0.567*** | 0.502*** | 0.572*** | 0.92*** | 0.891*** | 0.934*** | 0.313** |  |  | 0.402*** | 0.32* | 0.416** |
| FA | 0.315** | 0.521*** |  |  |  |  | 0.285* | 0.476** |  | 0.229* | 0.373* |  |  |  |  |  | 0.374* |  |  |  |  |  |  |  |
| GlcCer | 0.515*** | 0.617*** | 0.417** | 0.385*** | 0.341* | 0.478** | 0.504*** | 0.627*** | 0.407** |  |  |  |  |  |  |  |  |  |  |  |  |  |  |  |
| LPC |  |  |  |  |  | 0.308* |  |  |  |  |  |  |  |  |  |  |  |  |  |  |  |  |  |  |
| LPE | 0.258* | 0.33* |  |  |  |  |  | 0.304* |  |  |  | 0.343* |  |  |  |  |  |  |  |  |  |  |  |  |
| PC | 0.767*** | 0.792*** | 0.792*** | 0.36*** | 0.466** |  | 0.566*** | 0.641*** | 0.569*** | 0.507*** | 0.499*** | 0.683*** | 0.309** |  | 0.553*** | 0.519*** | 0.498*** | 0.705*** | 0.273* | 0.301* | 0.367* | 0.261* |  | 0.425** |
| PE | 0.606*** | 0.615*** | 0.614*** | 0.419*** | 0.662*** |  | 0.371*** | 0.399** | 0.378* | 0.424*** |  | 0.634*** |  |  | 0.401* | 0.446*** |  | 0.651*** | 0.25* |  | 0.381* | 0.304** |  | 0.471** |
| PI | 0.592*** | 0.495*** | 0.671*** |  | 0.406*** |  | 0.439*** | 0.334** | 0.519*** | 0.489*** | 0.393** | 0.574*** | 0.383*** |  | 0.562*** | 0.531*** | 0.392** | 0.626*** | 0.336*** | 0.387** | 0.317* | 0.386*** | 0.45*** | 0.376** |
| SM | 0.822*** | 0.825*** | 0.902*** | 0.313** | 0.415** |  | 0.758*** | 0.795*** | 0.831*** | 0.255* |  | 0.417** | 0.367*** | 0.32* | 0.633*** | 0.273* |  | 0.459** | 0.328** |  | 0.535*** |  |  | 0.367* |
| TG | 0.526*** | 0.524*** | 0.555*** | -0.309** |  | -0.337* | 0.369*** | 0.44** | 0.323* | 0.94*** | 0.949*** | 0.934*** | 0.64*** | 0.615*** | 0.624*** | 0.946*** | 0.95*** | 0.942*** | 0.317** |  |  | 0.437*** | 0.392** | 0.427** |

\*: p< 0.05; \*\*: p< 0.005, \*\*\*: p<0.0001 Acylcarnitine = 6 metabolites; Ceramide: 20, cholesteryl esters (CE): 28 metabolites; Diglycerides (DG) 4 metabolites; Fatty Acids (FA) 15; Glucosylceramide (GlcCer): 9 metabolites; Lysophosphatidylcholine (LPC) 29; Lysophosphatidyl ethanolamine (LPE) 4; Phosphatidylcholine (PC): 134; Phosphatidylethanolamine (PE) 31; Sphingomyelin (SM) 62; Triglycerides (TG) 83 metabolite

### Sex differences in correlation of CAPS and PCL-C scores with lipid subclasses

Next, we ascertained correlation of lipid subclasses with general health and PTSD symptom cluster measures (Table 1). Regression analysis revealed sex differences in correlation of 13 lipid subclasses with various PTSD, other biological and anthropomorphic measures. In sex aggregated analysis, 5/12 lipid subclasses associated with CAPS and PCL avoidance (C) scores, whereas only DGs associated with CAPS_C in women (r=0.38) and SM with CAPS_C in men (r=0.444). In women, only PE associated with PCL-C (r=0.36), whereas 7 subclasses associated significantly in men (Table 1). Interestingly, the same 7 lipid subclasses in men also associated with BMI, whereas in women, ceramide, DG, and TG associated with BMI. In men, LPE, and PE associated positively with age, whereas FA associated negatively, whereas no lipid subclass associated with age in women. Interestingly, in sex aggregated analysis, 5 lipid subclasses associated with body weight, but sex-specific analysis revealed that while DGs and TGs associated with body weight in women, no correlation was found in with lipid subclasses in men (Table 1).

### Integrated and systems lipidome and clinical measures analyses

The lipid panel identified a total of 413 known of which 348 were present in all 80 individuals; known metabolites were part of 13 classes of lipids namely, acylcarnitine, cholesteryl esters (CE), ceramides (Cer), glucosylceramide (GlcCer), diglycerides (DG), fatty acids (FA), LPC, lysophosphatidylethanolamine (LPE), PC, PE, PI, sphingomyelin (SM), and triglycerides (TG). Metabolomics in conjunction with clinical measures provides large datasets that are often difficult to interpret, thus an integrated and systems analysis approach is needed. PCA revealed that individuals with moderate and high PCL scores fell into discrete groups with women and men showing clear separation (Fig. 3a), suggesting sufficient variability in the datasets of men and women with PTSD, despite similar PCL scores and PTSD symptoms. Hence, all subsequent analysis was performed in a sex segregated manner and adjusted for BMI and PSQI. Volcano plot revealed that men with high PCL score experienced significant alterations in 8 lipid subclasses, whereas phosphatidylethanolamines were the only changed lipid subclass in women with high PCL score (Fig. 3b-d). PCA donut charts revealed subclasses of lipids that comprise various components to explain >90% variability in the lipidome in men with moderate and high PCL scores (Fig. 3c).

**Figure 3.**
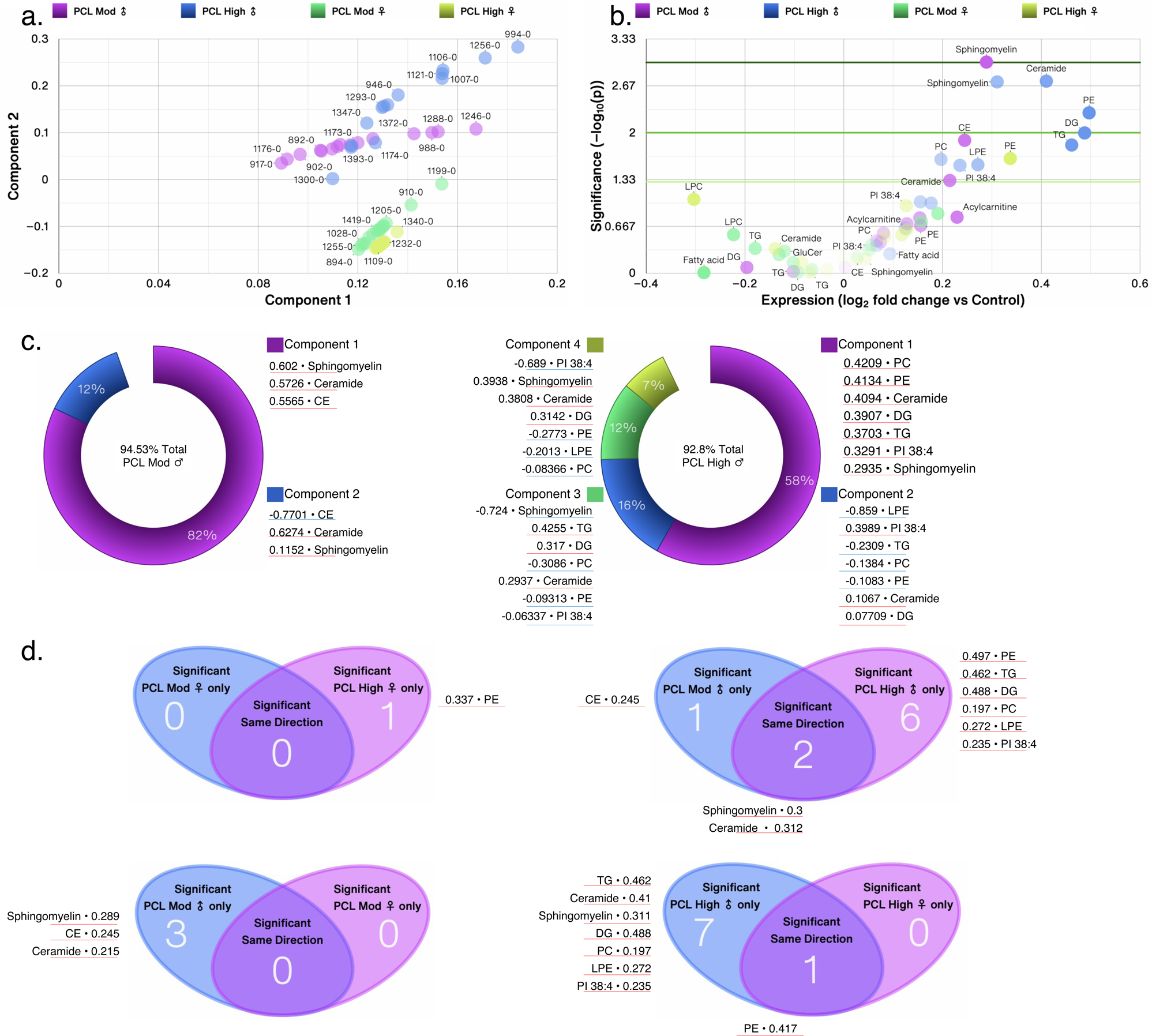
Sex differences in lipid subclasses in PTSD patients. (**a**) Principal component analysis (PCA) biplot showing clustering of samples by PCL scores and biological sex. (**b**) Volcano plots showing significantly changed 13 lipid subclasses in moderate and high PCL groups by sex compared with respective low PCL groups. The size of a point represents its quantifiability, calculated as 0.375 + 0.625 * sqrt([percent of cells positive]) * [percent of samples positive] * [maximum point size]; the opacity of points represents expression. The color intensity of points represents the size of the change, with increased symbols drawn in red and decreased symbols in blue. Green lines denote (from top to bottom) p < 0.001, 0.01 and 0.05 by Welch’s t-test. (**c**) PCA donut plots showing the primary components necessary to explain at least 90% of the variance in moderate (left) and high (right) PCL scores in male patients compared with low PCL scores for 13 lipid subclasses most correlated with each of the primary components versus low PCL scores of the same sex. No lipid subclass was significantly different in women with moderate PCL score and only phosphatidylethanolamine (PE) was significantly increased in women with high PCL scores compared with women with low PCL scores. (**d**) Venn diagram contrasting significantly changed measures between female (top left) and male (top right) PTSD patients each compared to non-PTSD patients and listing the most changed symbols for each Venn diagram segment, sorted by ascending p-value (not shown), with log_2_ fold change versus low PCL score non-PTSD patients shown. Venn diagram contrasting significantly changed measures between male and female PTSD patients with moderate PCL (bottom left) and high PCL (bottom right) scores. Lipid subclass measures with red lines were increased and those with blue lines were decreased.

### Men with high PCL score and severe PTSD have many more changes in lipidome than women with similar scores

Next, we performed integrated analysis with 348 individual lipids and 93 clinical measures and determined significantly changed individual lipid metabolites and clinical measures in women and men with moderate and high PCL scores versus women and men with low PCL scores, respectively. PCA donut charts revealed that component 1 explained 35% and 44% variability in the datasets (including lipidome and clinical measures) in women and men, respectively (Fig. 4a). Detailed distribution of first two PCA components is shown in scatter plots (Fig. 4b). Women with moderate and high PCL scores shared only ∼1.86% (8/430) measures; ∼3.2% (14/430) measures were significant only in PCL moderate group and ∼5.6% (24/430) measures were significantly altered in women with high PCL score compared with women with low PCL scores and no PTSD (Fig. 4c). In contrast to women, men with moderate and high PCL scores shared ∼10% (44/431) measures, with ∼4.2% (18/431) measures specifically altered in men with moderate PCL score and ∼21% (92/431) measures specifically altered in men with high PCL scores compared with men with low PCL scores and no PTSD (Fig. 4d). Pearson’s correlation of all clinical and lipid metabolites with each measure are shown in Supplementary Table S2. Men with moderate and high PCL scores demonstrated many more changes in individual lipid metabolites than women with similar PCL scores (Fig. 5a-b). Men with high PCL score and severe PTSD had 22% (94/431) significantly changed lipid metabolites, whereas women had only ∼5% (18/430) significantly changed measures with only 0.5% (3/431) measures (2 phosphatidylethanolamine (PE) and cholesterol) shared between women and men (Fig. 5c). When the analysis was performed including various clinical measures, the significantly shared measures increased to 8 from 5 (Fig. 5c, left).

**Figure 4.**
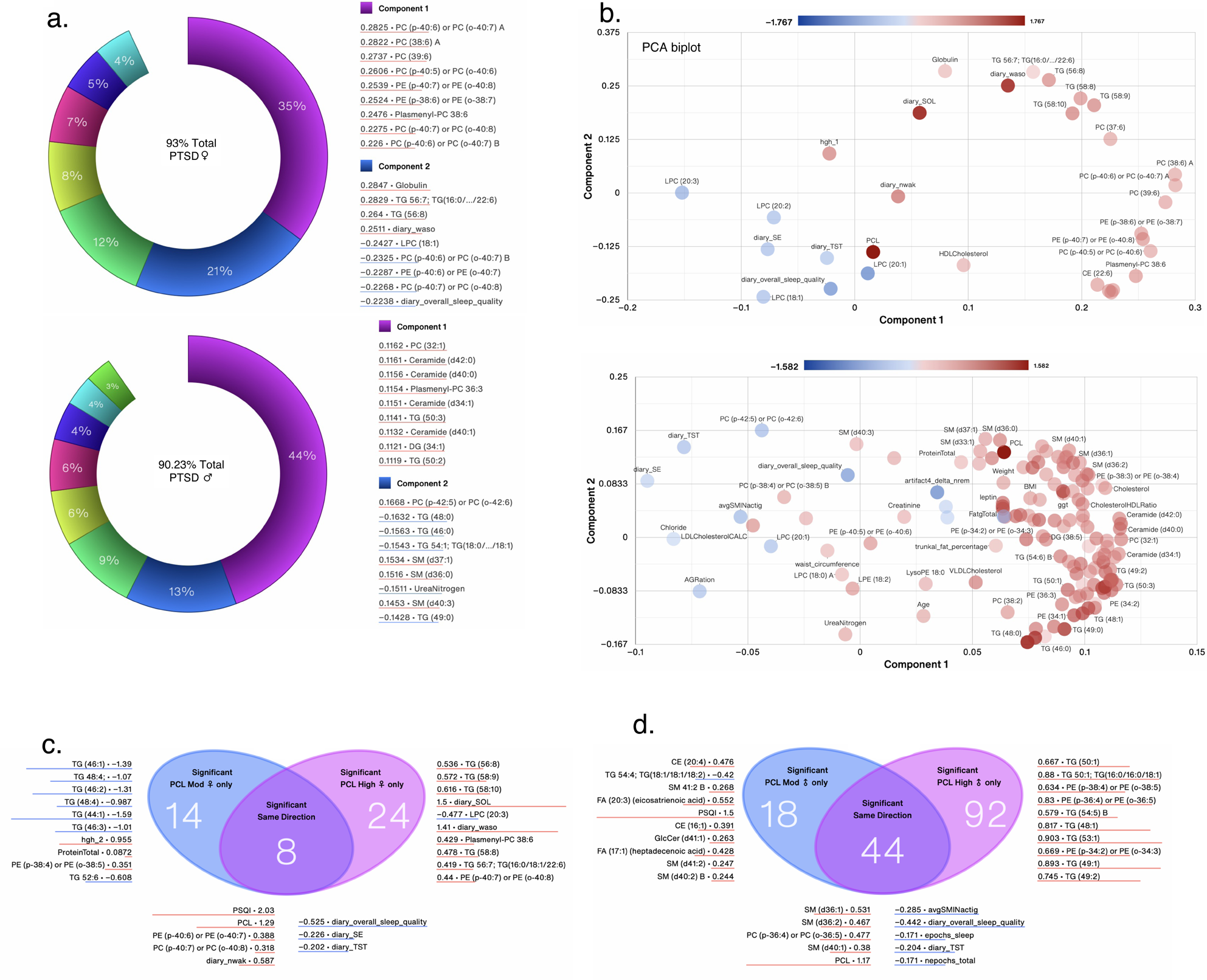
Integrated and systems lipidome and clinical measure analysis. (**a**) PCA donut plots showing the primary components necessary to explain at least 90% of the variance in females (top) and males (bottom) with high PCL scores compared with low PCL scores for all 348 individual lipid metabolites from 13 lipid subclasses and the 7 symbols most correlated with each of the first two primary components. (**b**) PCA biplots for the first two components with the color of points denoting log_2_ fold change versus low PCL scores of the same sex. PCA biplot showing clustering of samples by PCL scores and biological sex. Venn diagram contrasting significantly changed measures between female (**c**) and male (**d**) patients with moderate and high PCL scores compared to non-PTSD patients with low PCL scores and listing the most changed symbols for each Venn diagram segment, sorted by ascending p-value (not shown), with log_2_ fold change versus low PCL score non-PTSD patients shown. Lipid metabolites and clinical measures with red lines were increased and those with blue lines were decreased.

**Figure 5.**
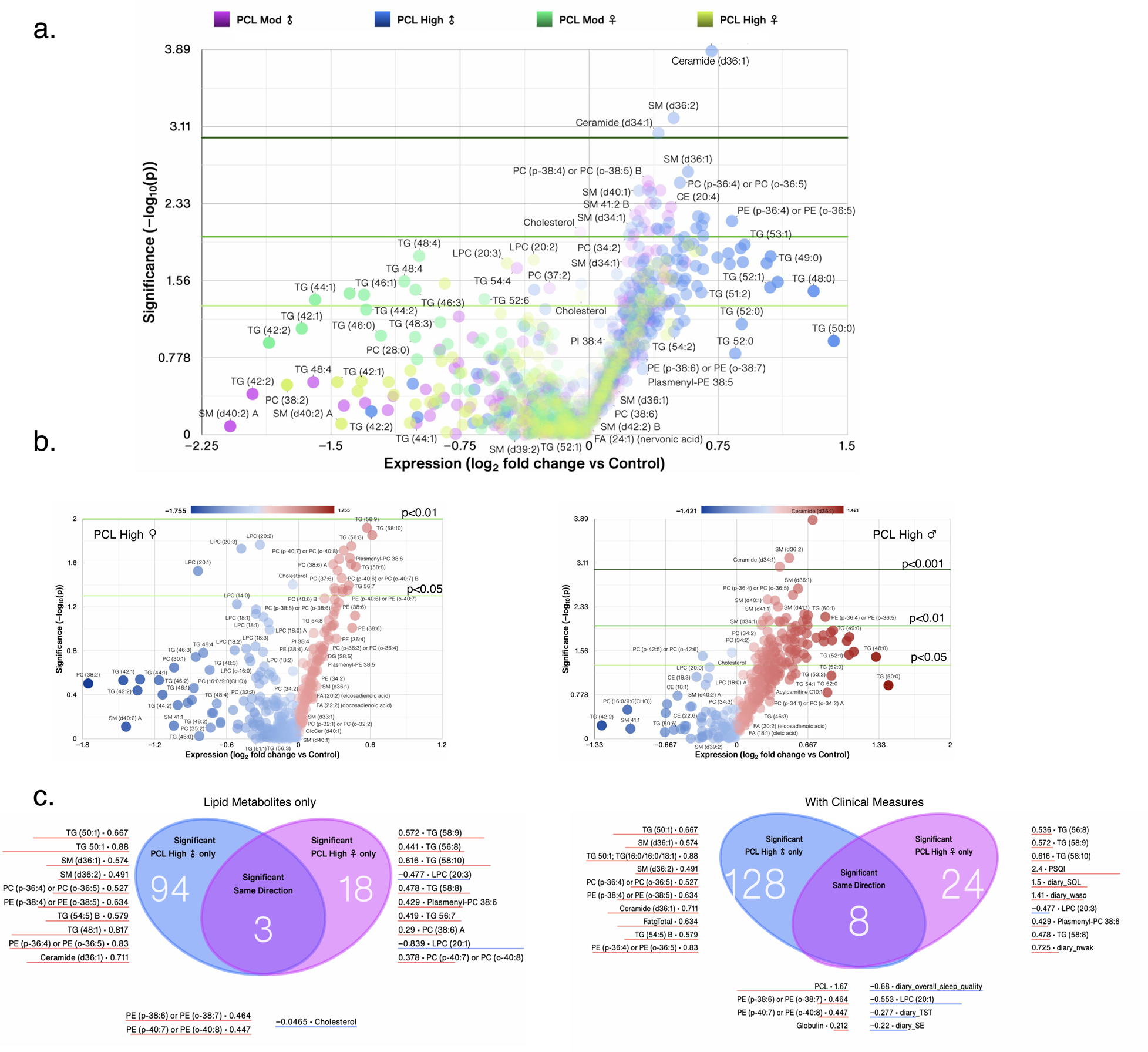
Sex differences in lipid metabolites in PTSD patients. Venn diagram contrasting significantly changed lipid metabolite measures for (**a**) all subgroups, (**b**) female and male patients with high PCL scores compared to non-PTSD patients with low PCL scores. The size of a point represents its quantifiability, calculated as 0.375 + 0.625 * sqrt([percent of cells positive]) * [percent of samples positive] * [maximum point size]; the opacity of points represents expression. (**c**) Venn diagram contrasting significantly changed lipid measures only (left) or lipid and clinical measures combined (right) between female and male patients with high PCL scores and listing the most changed symbols for each Venn diagram segment, sorted by ascending p-value (not shown), with log_2_ fold change versus low PCL score non-PTSD patients shown. log_2_ fold change was calculated for each sample versus its corresponding value with low PCL score, with the average shown and *, ** and *** denoting p < 0.05, 0.01 and 0.001 by Welch’s t-test, respectively. Lipid metabolites and clinical measures with red lines were increased and those with blue lines were decreased.

### Sex differences in clinical and sleep measures in individuals with PTSD

Health conditions such as obesity contribute to several changes in clinical outcomes including changes in heart, liver function as well as disturbed sleep. We next determined what clinical laboratory values as well as sleep measures differed between men and women with PTSD compared with non-PTSD individuals. Using heat maps, we compared side-by-side, all significantly changed clinical and sleep measures in men and women with moderate and high PCL scores compared with men and women with low PCL scores, respectively. BMI, age, total fat, cholesterol, triglycerides, creatinine, leptin, GGT were among some of the clinical measures (24/97) that were significantly increased in men, but not women (Fig. 6a). GGT levels were increased ∼2-fold in men with high PCL score (log_2_ fold=1), whereas GGT levels were increased in women with high PCL scores but did not reach statistical significance (Fig. 6a). Total protein, human growth hormone, calcium, albumin, and HDL cholesterol levels were higher in women with moderate and high PCL scores compared with women with low PCL (Fig. 6a). Both women and men with moderate-to-high PCL scores reported worst perceived sleep quality with significantly worse PSQI scores (Fig. 6b). Several other measures of sleep, both objective and qualitative, such as in_delta_nrem, total epochs, and AGRation were significantly decreased in men with high PCL scores compared with men with low PCL scores (Fig. 6b). Women with high PCL scores reported significantly more waking after sleep onset (WASO) compared with women with low PCL scores and men with PTSD, regardless of severity (Fig. 6b). Women with moderate PCL scores experienced changes in only 10/430 lipid metabolites, 6 triglycerides and 1 PC (40:6 A) were decreased and 3 PE/PC were increased compared with women with low PCL scores (Fig. 6c). Women with high PCL scores experienced changes in 20/430 lipid metabolites with increases in TG, PC, PE, and CE subclasses, whereas LPCs decreased (Fig. 6d).

**Figure 6.**
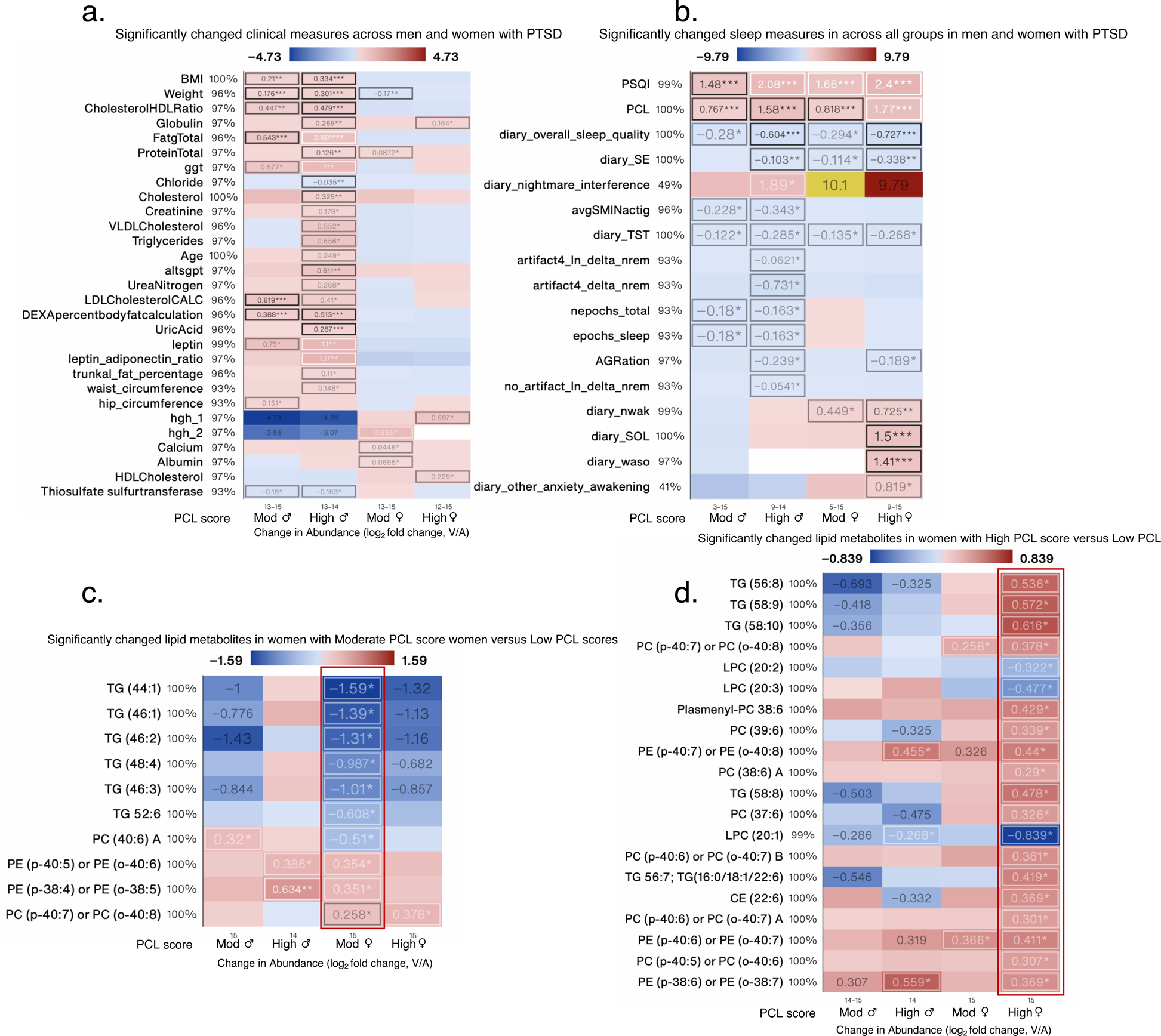
Significantly changed clinical measure in all PTSD patients and lipid measures in female PTSD patients. Heat map of the most-significantly changed clinical (**a**), sleep (**b**) measures in men and women with high PCL scores, lipid metabolite measure in women with moderate PCL score (**c**) and high PCL score (**d**) compared with men and women with low PCL scores, respectively. Changed measures are shown for all 4 subgroups. log_2_ fold change was calculated for each metabolite versus its corresponding metabolite value in low PCL group, with the average shown and *, ** and *** denoting p < 0.05, 0.01 and 0.001 by Welch’s t-test, respectively. Labels next to symbols denote the percentage of samples in which metabolites were detected across all groups; group sizes (n) are shown above group names.

In men with moderate PCL score, 24/62 sphingomyelins (SM) were increased and were the most changed subclass of lipids (Fig. 7a). Other lipid metabolites increased were from subclasses acylcarnitine, ceramide, CEs, GlcCer, fatty acids, PCs, and LPC (Fig. 7b). In men with high PCL score, PE (18/31), PC (14/134), LPC (3/29), SM (20/62), TG (18/34), and ceramide (14/20) along with a select number of fatty acids, acylcarnitine, LPE, PI, and DG were also changed (Fig. 7c-f). Interestingly, several SM, TG and Ceramides that were increased in men with high PCL score, trended to decrease in women with moderate-to-high PCL score, suggesting divergent responses to similar trauma.

**Figure 7.**
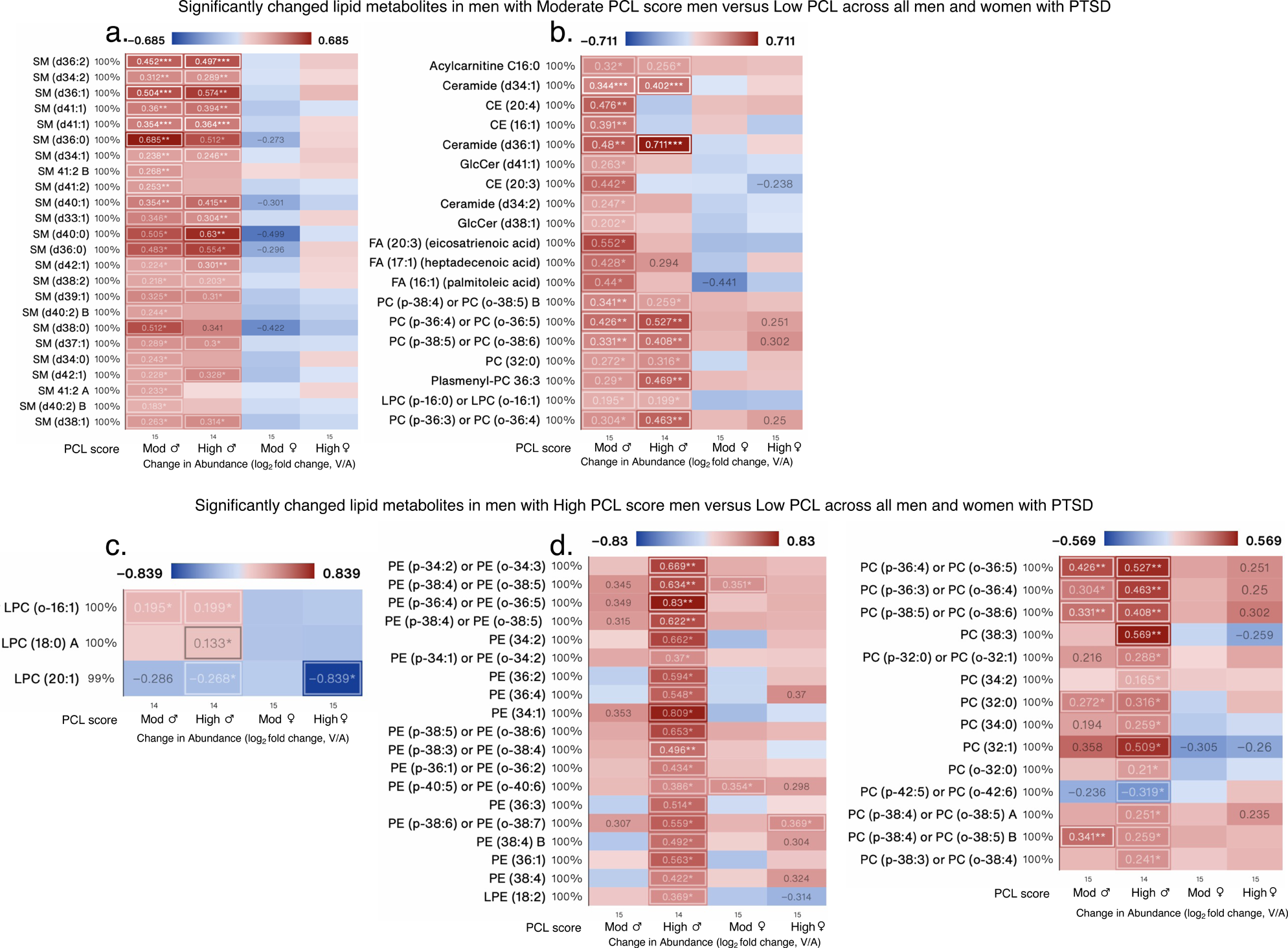
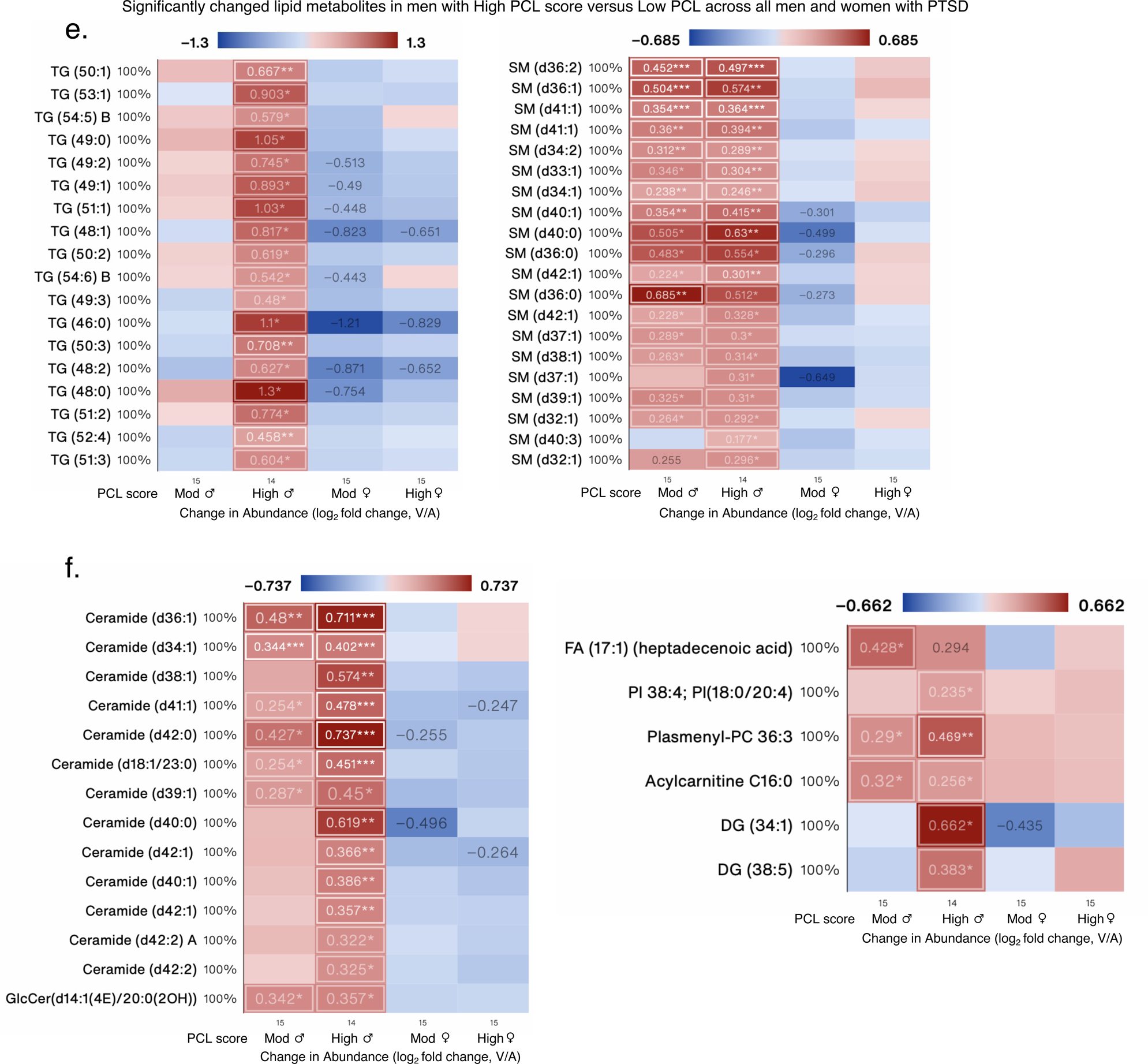
Significantly changed lipid measures in male PTSD patients. Heat map of the most-significantly changed lipid measures in (**a-b**) men with moderate PCL scores, and (**c-f**) men with high PCL scores compared with men with low PCL scores, respectively. Changed measures are shown for all 4 subgroups. Many of the lipid metabolites increased in men with high PCL score were decreased in women. Log_2_ fold change was calculated for each metabolite versus its corresponding metabolite value in low PCL group, with the average shown and *, ** and *** denoting p < 0.05, 0.01 and 0.001 by Welch’s t-test, respectively. Labels next to symbols denote the percentage of samples in which metabolites were detected across all groups; group sizes (n) are shown above group names.

### Most changed individual metabolites in individuals with moderate and high PCL scores correlate with cholesterol and other lipids, but not PCL or PSQI

We next determined correlation between top changed individual lipid metabolites in individuals with PTSD with clinical measures and other lipid metabolites. TG (44:1) was the most changed/decreased (log_2_ fold=-1.59, p <0.05) in women with moderate PCL score versus women with low PCL score (Fig. 6c). TG (44:1) correlated most significantly in all groups with LPC (14:0) (r-0.665, p=10^−8.2^); however, the correlation was stronger in men with high PCL score than in women (Fig. 8a, ♂: r=0.855, p=10^−4^ vs ♀: r=0.684, p=0.0049, respectively). TG (44:1) also correlated with VLDL cholesterol in men and women with high PCL, but not low PCL scores (Fig. 8a, ♂: r=0.701, p=0.007 and♀: r=0.533, p=0.04). TG (56:8) was one of the top and most significantly increased metabolites in women with high PCL score (log_2_ fold= 0.536, p <0.05, Fig. 6D) and it correlated with VLDL cholesterol in women, but not men with high PCL scores (Fig. 8b, ♀: r=0.821, p=10^−3.8^ and ♂: r=0.407, p=0.167). PC (40:6)B was the most correlated lipid metabolite with TG (56:8) in all patients (r=0.664, p=10^−8.1^), however, individuals with low PCL scores displayed a very strong and robust correlation with a weaker correlation in women with high PCL score (Fig. 8b). In men with moderate and high PCL scores shared many of the changed metabolites. Ceramide (d40:0), Ceramide (d36:1), SM (d40:0), and TG (48:0), were amongst the most increased metabolites in men with high PCL scores (Fig. 7e-f). All the metabolites correlated with some form of cholesterol (Fig. 8c-d) and TG (48:0) also correlated robustly in all patients with total sleep time and systolic blood pressure (Fig. 8d, r=0.624, p=10^−7^, and r=0.42, p=10^−3.1,^ respectively). However, the correlation mostly held in men with high PCL scores and not in women (Fig. 8d).

**Figure 8.**
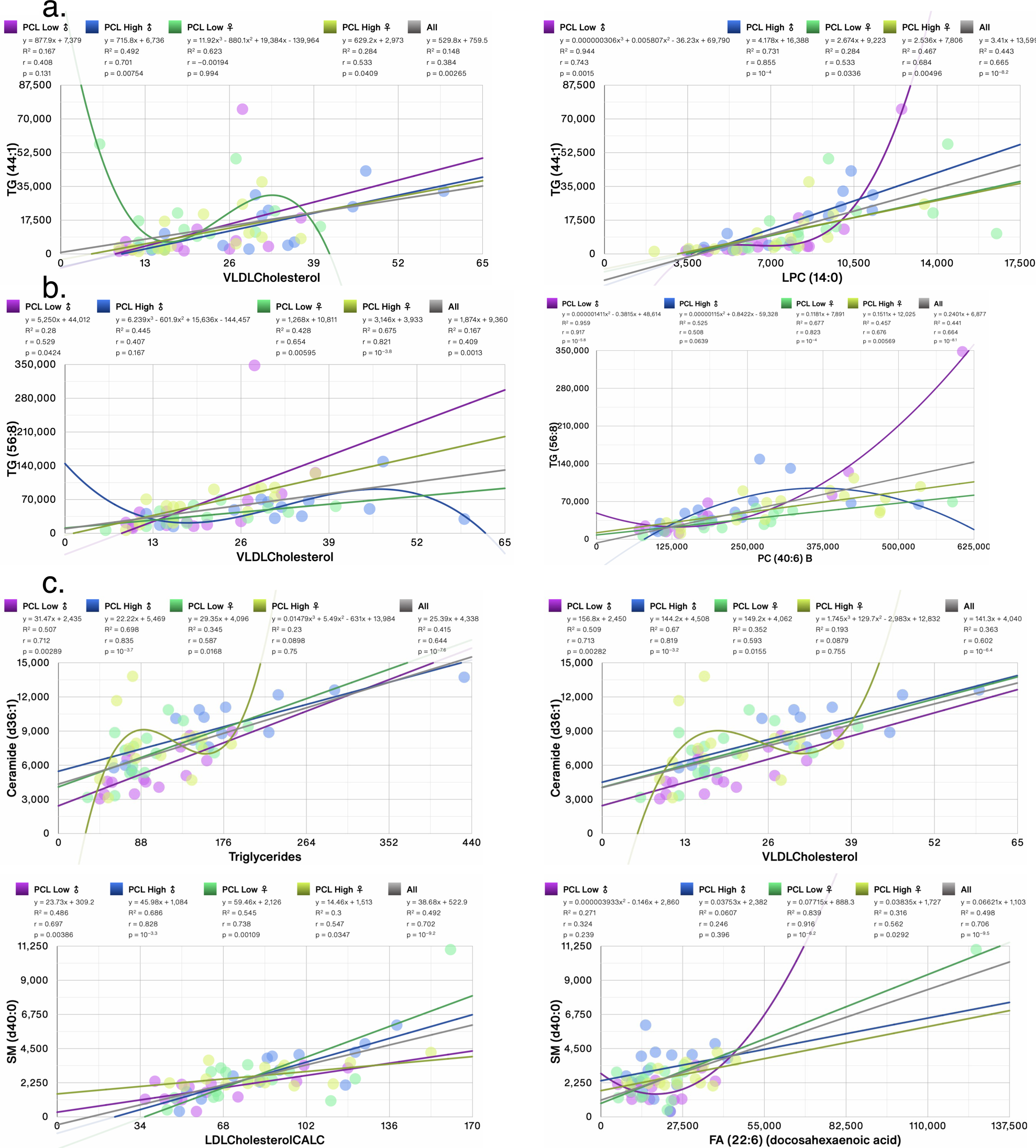
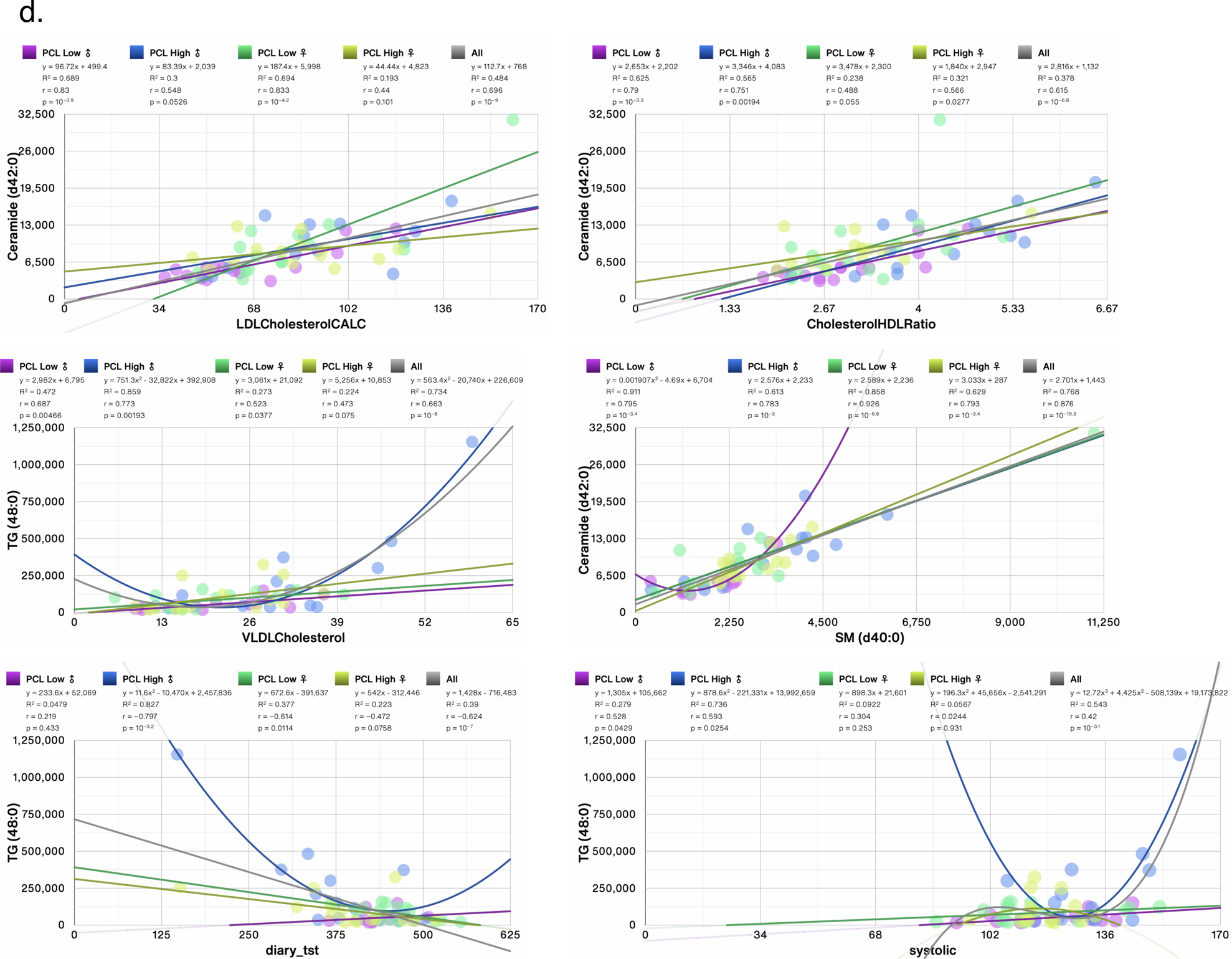
Scatter plots showing correlations between top 2 most significantly changed lipid metabolites in (**a**) females with moderate PCL score (TG (44:1)), (**b**) females with high PCL score (TG (56:8)), (**c**) males with moderate PCL score (Ceramide (d36:1) and SM (d40:0)), (**d**) males with high PCL score (Ceramide (d42:0) and TG (48:0)). Trendlines, or an n^th^-degree polynomial trendline if its goodness-of-fit is either 50% greater than, or if it explains at least half of the variance not explained by the (n – 1)^th^-degree polynomial, were fit to the data. R^2^, r and p denote goodness-of-fit, Pearson’s correlation coefficient and significance of the correlation, respectively. Several changed metabolites correlated with clinical measures of (V)LDL cholesterol. Measures also correlated with triglycerides, systolic blood pressure, and total sleep time (tst). Most significant correlations with other lipid metabolites are also shown.

## Discussion

Few studies have examined lipid metabolites in PTSD. Most of the cohorts studied consisted primarily of males and none have analyzed data in a sex segregated manner^29,32–34^. We have previously examined primary metabolites in male and female PTSD patients and reported sex differences in levels of several metabolites and the contribution of perceived sleep quality in many of the changes in primary amines^14^. No study thus far has integrated clinical health measures with lipid metabolites or analyzed the contribution of comorbidities such as sleep disorders in the altered lipid profile. Here, we performed, to our knowledge, first known integrated and systems levels analysis of clinical health, sleep measures and lipid metabolites measured using QToF mass spectrometry in individuals with PTSD and trauma-exposed non-PTSD individuals. Many more lipid metabolites were altered in men with higher PTSD symptoms than in women with similar scores. Many lipid metabolites that were significantly increased in men with PTSD compared with non-PTSD men belonged to lipid subclasses that included TG, SM, Ceramide, and PE; several of these metabolites were decreased in women with PTSD, although the decreases were not significant from non-PTSD women. The use of metabolomics to better understand the pathophysiology of PTSD has great potential since it considers all known, and even unknown, molecules and pathways all at once. Metabolomics is a global and unbiased approach to understanding regulation of metabolic pathways and networks of physiologically relevant interactions. The metabolome is regulated by gene-environment interactions and reflects the intermediary state between genotype and phenotype. Technological developments such as mass spectrometry allow for the examination of metabolites and discovery of novel pathways using an unbiased method to examine multiple analytes simultaneously and hence offers a great advantage to detect patterns of alterations in a complex disorder such as PTSD.

Several studies have examined the lipidome of PTSD. One study examined peripheral blood serum samples of 20 PTSD civilian patients and 18 healthy non-trauma controls^35^ and found alterations in lipid-derived metabolites were altered in PTSD. Unfortunately, sex differences were not able to be examined due to the small sample size. In contrast to these findings, PTSD was associated with alterations in lactate and pyruvate, pathways related to glycolysis, decreases in unsaturated fatty acids involved in inflammatory processes, and metabolism, possibly pointing to mitochondrial alterations in male combat trauma-exposed veterans from the Iraq and Afghanistan conflicts (n=52 PTSD; n=51 controls). Two glycerophospholipids, phosphatidylethanolamine (18:1/0:0) and phosphatidylcholine (18:1/0:0) ^36^ that may relate inflammation, mitochondrial dysfunction, membrane breakdown, oxidative stress, and neurotoxicity were found in a study of male Croatian war veterans (n= 50 with PTSD, 50 healthy controls; and a validation group of n=52 with PTSD, n=52 healthy controls). Alterations in compounds related to bile acid metabolism, fatty acid metabolism and pregnenolone steroids, which are involved in innate immunity, inflammatory process and neuronal excitability were found in World Trade Center male responders (n=56 PTSD, n=68 control)^37^. The largest study to date examined differences in severity and progression of PTSD using a multi-omics analysis of metabolomics, proteomics and DNA methylome assays in 159 active-duty male participants with relatively recent onset PTSD (<1.5 years) and 300 male veterans with chronic PTSD (>7 years)^38^. Their findings indicated that active-duty participants with recent PTSD had alterations in signaling and metabolic pathways involved in cellular remodeling, neurogenesis, molecular safeguards against oxidative stress, polyunsaturated fatty acid metabolism, immune regulation, post-transcriptional regulation, cellular maintenance, and markers of longevity. In contrast, Veterans with chronic PTSD showed evidence of alterations associated with chronic inflammation, neurodegeneration, cardiovascular and metabolic disorders, and cellular attrition. These findings suggest that time since trauma and/or age may play an important role, as molecular alterations in the younger cohort reflected homeostatic or compensatory responses, whereas the older cohort had alterations indicative of more chronic disease. The implications of these findings may be relevant for women in the menopausal transition; however, too few women were included in this latter study to allow for an analysis of sex differences.

One of the most common complaints among individuals with PTSD is sleep disturbance. Lower slow wave sleep duration and delta-band spectral power are more pronounced in men than women^3^, and correlate with PTSD status In contrast, greater rapid eye movement sleep is found in women with PTSD compared to healthy controls, a difference not seen in men^3^. In this study, we found that WASO, minutes of wake that occurs between sleep onset and waking was over ∼2.5-fold greater in women with severe PTSD symptoms compared with no PTSD or with men with PTSD. Sleep disturbance is a key risk factor for health consequences as it alters hypothalamic-pituitary-adrenal (HPA) axis function, resulting in impaired glucose and lipid metabolism. The HPA and somatotropic axes activities are temporally associated with delta power sleep and promote insulin sensitivity and metabolic syndrome. Additionally, sleep duration correlates with metabolic risk in PTSD but does not fully account for the association between PTSD with known metabolic disturbances such as in blood insulin or glucose levels^39^. We have previously shown opposite correlations between tryptophan and insulin levels in men and women with PTSD^14^. Moreover, while estrogen, progesterone, and testosterone and their metabolite levels did not differ between individuals with and without PTSD in this cohort, delta power sleep correlated with testosterone levels in men^14^. In this cohort, among men, one triglyceride metabolite (TG (48:0)) was negatively correlated with total sleep time with greater PTSD severity and systolic blood pressure. Among women, TG (48:0) was correlated with lower PTSD severity. Men in this cohort with high PTSD severity also had higher BMI and the correlation between BMI and systolic blood pressure were inversely correlated in men but not women (PCL low ♂: r=0.701, p=0.003 vs PCL High ♂: r= −0.212, p=0.466). BMI was no longer correlated with triglycerides in men (PCL low ♂: r=0.614, p=0.014 vs PCL High ♂: r= 0.498, p=0.07) or women with PTSD. Thus, higher weight might be contributing to changes in metabolic health in men with PTSD as women with PTSD in this cohort did not have significant changes in metabolic health and had BMI within normal range. Others have reported associations between PTSD symptoms and three specific glycerophospholipids in Veterans with PTSD^32^; these specific metabolites were not ascertained in our cohort. However, no information on BMI and other clinical measures was reported or is available for comparison.

Sphingomyelins regulate endocytic function as well as receptor-mediated ligand uptake of ion channels and G protein-coupled receptors^40^. SMs also regulate cardiovascular function and their distribution within cellular compartments often correlates with cholesterol. We found nearly 65% of SM (24/37) were increased in men with moderate and severe PTSD compared with low/no PTSD, whereas the same SM were decreased in women with moderate PTSD and either increased or decreased in women with severe PTSD compared with women with no PTSD, however, the differences in women did not attain statistical significance. Sphingomyelins (>40 carbon length) were reportedly increased in a cohort of World Trade Center responders with PTSD compared with non-PTSD individuals who were also exposed to the 9/11/2001 attacks in New York City^34^. In the World Trade Center cohort, after adjusting for several medical conditions, 9 different SM correlated with BMI in their cohort, whereas hexosylceramide (HCER (26:1) did not. Kuan et al. also performed integrated analysis of lipid metabolites with proteomics and found several protein modules that contained IL-6 and ATP6V1F proteins associated with fatty and bile acid metabolites^34^. However, Kuan et al. could not replicate findings from other 3 studies ^29,32,33^ about specific lipid metabolites and their correlations with PTSD status, likely due to several variables, including clinical profiles and PTSD measures used^34^. Our aim was not to replicate findings but perform an unbiased analysis. Adjusting for BMI, PSQI or other clinical measures (such as total protein, cholesterol etc.) or not, did not alter our lipid metabolite findings.

Many essential fatty acids are derived from diet. Diet is one variable that is difficult to control for and is a limitation for metabolomics analysis. Individuals in our cohort were in a 3-day/night sleep clinical and had access to limited dietary choices. Thus, while our cohort is not diet-controlled, per se, dietary variability is far less than those that might be present in other studies that have interrogated the lipidome of individuals with PTSD. In this regard, we identified over 1500 unknown lipid metabolites. Many of these were highly altered in individuals with PTSD, more than any of the known lipids. In the future, identification of these unknown metabolites can lead to discovery of PTSD-specific lipid markers. While medical conditions may contribute to alterations in lipidomics, this sample included young and healthy participants and there were no associations between PTSD and chronic disease states in this sample. It is possible that alterations in metabolites may reflect earlier stages of future disease. Contrary to our expectation that individual lipid metabolites that were changed in men and women with moderate and high PCL scores should correlate to some degree with PCL scores and sleep quality; we did not find any. While prior studies have found alterations in neuroendocrine, immune, and aging processes in PTSD, our knowledge of the role of metabolite disturbances is PTSD is limited but has the potential to elucidate new discovery of yet unknown biological mechanisms of disease.

While highly promising, there are several limitations to the metabolomics research to date: 1) While these studies point to several plausible biological pathways with the potential to explain alterations in PTSD and common comorbidities, a major limitation is that these studies were conducted almost entirely in males or had insufficient numbers of female participants to examine sex differences. Indeed, some of the differences in findings may be attributed to differences in sex as there were differences in findings in the studies that included women, but this has not been tested. 2) The studies that included women did not control for menstrual cycle day. Since sex steroids may interact with metabolites in other biological pathways and may fluctuate over the menstrual cycle, determining menstrual cycle day and examining sex steroids along with other metabolites of interest is critical. 3) All studies used cross-sectional designs with single time point blood draws. Some control groups were not trauma exposed. Thus, it is unclear if these findings reflect PTSD symptom state, trait, or a response to trauma exposure regardless of the presence of PTSD symptoms. To establish biomarkers of disease, it will be important to determine whether these alterations are stable over repeated measures. 4) A major challenge to finding a reliable biological signal in PTSD lies in the heterogeneity of the disorder. There is a great deal of variability in the symptom profile and in combinations of symptoms in PTSD, as described by Galatzer-Levy^20^. PTSD symptoms are typically assessed by self-report or diagnostic interview, resulting in subjective perceptions of severity which can be unreliable. Given that PTSD is a heterogenous disorder, analysis by PCL score, rather than grouping all people with PTSD diagnosis with large variation in scores, yields more robust findings. There is also substantial comorbidity between PTSD and other mental health conditions such as depression, traumatic brain injury or substance use, among others. Overlapping symptoms may obscure biological findings.

Our study has several strengths-first, we performed an unbiased analysis of lipids using validated QToF mass spectrometry. Second, we performed an integrated and systems level analysis taking clinical measures into account. Third, in-depth sleep measures were obtained. Fourth, the non-PTSD population was age-matched and trauma-exposed. Caveats include a cross-sectional study design. A longitudinal study design would be needed to demonstrate stability of metabolite changes over time over the course of the disease process. In our ongoing studies, longitudinal sample collection from PTSD patients is ongoing and data from that study should address some of these limitations in the future.

## Materials and methods

### Human subjects

This study was a 2 x 2 cross-sectional design with 4 groups (PTSD/control × women/men) of 44 individuals with current chronic PTSD and 46 control subjects. Participant’s age ranged from 19-39 years. Data from 4 participants were excluded due to difficulties in blood collection. Sleep measures were recorded in an inpatient sleep laboratory at the General Clinical Research Center (GCRC) at the University of California, San Francisco. The Committee on Human Research at the University of California, San Francisco approved this study. Written informed consent was provided to all participants before enrollment and start of any study procedures.

PTSD subjects met DSM-IV criteria as ascertained using the Clinician-Administered PTSD Scale (CAPS) or score >40 and as described earlier by us^3^. Control subjects had no lifetime or current history of a PTSD diagnosis. Women participants were premenopausal and were scheduled during the follicular phase of the menstrual cycle. All study procedures were timed according to habitual sleep onset, determined by actigraphy and sleep diary in the week prior to the GCRC study. Study participants were limited to one cup of caffeine daily, maintained regular bed and waking times, did not consume alcohol, and were drug-free. Exclusion criteria for PTSD and control subjects was as described previously^3^: history of traumatic brain injury, presence of neurologic disorders or systemic illness; use of psychiatric, anticonvulsant, antihypertensive, sympathomimetic, steroidal, statin or other prescription medications; obesity (defined as body mass index (BMI) >30); alcohol abuse or dependence in the prior two years; substance abuse or dependence in the previous year; any psychiatric disorder with psychotic features; bipolar disorder or obsessive-compulsive disorder; and pregnancy. Exclusion criteria for control subjects also included a lifetime history of major depressive disorder or panic disorder.

### Psychiatric diagnoses and trauma history

The Life Stressor Checklist-Revised interview was used to determine trauma exposure and age of occurrence^41^. The PTSD Checklist (civilian version) for DSM-IV (PCL-C)^42^ was used as a self-reported measure of severity of chronic PTSD symptoms^43^. The PCL-C consists of 17 items that correspond to the DSM-IV criteria and include intrusive thoughts and re-experiencing symptoms (cluster B), avoidance (cluster C), and hyperarousal (cluster D). All other psychiatric disorders, including major depressive disorder were diagnosed using the structured clinical interview for DSM-IV, non-patient edition (SCID-NP)^44^.

### Sleep clinic and measures

A subjective assessment of sleep quality, sleep latency, sleep duration, sleep efficiency, sleep disturbances (including nightmares) was obtained using the Pittsburgh Sleep Quality Index^45^ (PSQI). The use of sleep medication, and daytime dysfunction over the previous month has been described elsewhere^3^.

### Polysomnographic measurements and power spectral analysis for polysomnographic measures

Electroencephalogram (EEG) from leads C3, C4, O1 and O2, left and right electrooculograms (EOG), submental electromyogram, bilateral anterior tibialis EMGs, and electrocardiogram were recorded in accordance with standardized guidelines by Rechtschaffen^46^ using the Ambulatory polysomnography (Nihon Kohden Trackit Ambulatory Recording System) as described previously^3^. Sleep activity in all frequency bands delta through gamma from the C3 electrode was measured by power spectral analysis using the Pass Plus (Delta Software) analytic software. Delta sleep spectral power density (μV2) was natural log transformed to normalize its distributions.

### Blood collection and clinical laboratory measures

Blood was collected at habitual wake-up time on the morning after the second night on the GCRC, while the subject was fasting, from an indwelling catheter inserted the night before. Blood (10mL) was drawn into a chilled EDTA tube and processed for plasma separation for mass spectrometry analysis.

### Analysis of lipid metabolites in human plasma using liquid chromatography-Quadrupole Time-of-Flight mass spectrometry (LC-MS/MS)

Lipid metabolites that included diverse classes of phospholipids, ceramides, sphingomyelins, free fatty acids, acylcarnitines, triacylglycerides, cholesterol and cholesterol esters were measured using validated LC-MS/MS procedures^47^. Briefly, lipids from 40 µL plasma were extracted using 300 µL degassed, −20°C cold methanol 300 μL containing a mixture of lipid internal standards, specifically LPE(17:1), LPC(17:0), PC(12:0/13:0), PE(17:0/17:0), PG(17:0/17:0), d7-cholesterol, SM(d18:1/17:0), Cer(d18:1/17:0), sphingosine(d17:1), DG(12:0/12:0), DG(18:1/2:0), and d5-TG-(17:0/17:1/17:0). After adding 1 mL of cold methyl tertbutyl ether (MTBE) was added containing CE(22:1) as additional internal standard, lipids were separated from hydrophilic metabolites by adding 250 μL LC−MS grade water. Dried lipid extracts were resuspended in 110 µL methanol/toluene (9:1, v/v) containing CUDA as system suitability internal standard prior to LC−MS/MS analysis. 1.7 µL were injected onto an Acquity UPLC CSH C18 column (100 × 2.1 mm; 1.7 μm) coupled to an Acquity UPLC CSH C18 VanGuard precolumn (5 × 2.1 mm; 1.7 μm) (Waters, Milford, MA). The column was maintained at 65 °C at a flow-rate of 0.6 mL/min with a water/acetonitrile/isopropanol gradient using mobile phases (A) 60:40 (v/v) acetonitrile:water and (B) 90:10 (v/v) isopropanol:acetonitrile^48^. Both mobile phases were buffered with ammonium formate (10 mM) and formic acid (0.1%). Lipid separation was performed with the following gradient: 0 min 15% (B); 0−2 min 30% (B); 2−2.5 min 48% (B); 2.5−11 min 82% (B); 11−11.5 min 99% (B); 11.5−12 min 99% (B); 12−12.1 min 15% (B); and 12.1−15 min 15% (B). Mass spectrometry was performed on an Agilent 6530 quadrupole/time-of-flight mass spectrometer (QTOF MS) with a Dual Spray ESI ion source (Agilent Technologies, Santa Clara, CA). Simultaneous MS1 and data dependent MS/MS acquisition was used. Electrospray (ESI) parameters were set as: capillary voltage, 3.5 kV; nozzle voltage, 1 kV; gas temperature, 325 °C; drying gas (nitrogen), 8 L/min; nebulizer gas (nitrogen), 35 psi; sheath gas temperature, 350 °C; sheath gas flow (nitrogen), 11 L/min; MS1 acquisition speed, 2 spectra/s; MS1 mass range, m/z 60–1700; MS/MS acquisition speed, 2 spectra/s; MS/MS mass range, m/z 60–1700; collision energy, 25 eV. The instrument was tuned using an Agilent tune mix. A reference solution (m/z 121.0509, m/z 922.0098) was used to correct small mass drifts during the acquisition. For quality control (QC), we randomized injection orders, used pool QC samples to equilibrate the LC−MS system before data acquisition, injected method blank samples and a pool QC samples between each set of 10 study samples, and injected the NIST SRM 1950 community plasma QC sample before and after the study sample sequence. We also monitored peak shapes and intensities of all internal standards during data acquisition. Data were processed by MS-DIAL (v. 2.69) software program with MS1 (centroiding) tolerance 10 mDa, mass slice width 50 mDa, smoothing level 3 scans, minimum peak height 500 amplitude, and alignment at 25 mDa and 6 s retention time tolerance. Lipid were identified at 9 s retention time tolerance with accurate mass and MS/MS matching against the LipidBlast library^49^. Quantification was performed by combining the two most abundant adducts per lipid^50^, followed by normalization using the sum of all internal standards. Due to recursive backfilling during data processing, missing values were limited to 4.1% of all data. Such missing values imputed using half of the minimum detected value for each compound. All metabolomic data were investigated using pooled quality control samples and blank samples. Metabolites that had more than 30% relative standard deviation in pooled QC samples were removed, and metabolites that showed less than 3-fold higher intensity compared to blank samples were removed as well. Technical errors reduce biological power, so findings reported in this work were observed at statistical power despite possible technical variance.

### Statistical analysis

Aseesa’ Stars version 0.1 (www.aseesa.com) analysis tool was used for generation of heat maps, bar charts, principal component analysis, correlation scatter plots, volcano plots and Venn diagrams as described previously^51^ and below:

#### Heat Maps

Heat maps were generated to depict several symbols in a single chart instead of multiple bar charts as described in detail elsewhere^51^. Value labels show the group average or average log_2_ fold change vs the respective control group. Value labels are drawn for every symbol in absolute heat maps, and for values greater than 33% of the heat map’s maximum value in relative heat maps. Labels for values less than 16.67% of the maximum are drawn in black for legibility as reported earlier^51^. Labels next to gene symbols denote the percentage of participants across all test groups in which metabolite of the symbol were detected. Numbers above test group labels denote the number of samples that were included (n); if not all metabolites were present, then minimum sample size is shown.

#### Bar Charts and Volcano Plots

Values in bar charts are calculated in the same way as those in heat maps. Bar chart legends includes top four correlated measures sorted by significance value. Error bars represent the standard deviation. The comparison mode Value-to-Average was used as described previously^51^. Measures with p < 0.1 are included in volcano plots for clarity. Welch’s t-test was performed as in heat maps, with ^†^, ^††^ and ^†††^ denoting p < 0.05, 0.01 and 0.001 versus the previous test group (one bar above). Labels next to symbols denote the number of samples that were included (n). The filled fraction of a bar represents the percentage of cells in which transcripts of the symbol were detected; in charts showing changes in cell types, it represents the percentage of samples in which cells of that type were detected.

#### Principal Component Analyses (PCA)

For Symbol PCAs, only symbols with p < 0.05 were included, and samples with zero values were excluded. Covariance matrixes were created by standardizing all values for each symbol using 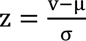 for a sample value v, group average μ and standard deviation σ, and calculating the covariance between two symbols^51^. Donut charts show the primary components necessary to explain at least 90% of the dataset’s variance. PCA biplots show the correlation of each included symbol/sample with Component 1 (x axis) and Component 2 (y axis) as given by the components’ eigenvectors. The color of points in biplots for Symbol PCAs denotes log_2_ fold change versus control, calculated in the same way as in bar charts, with increased symbols shown in red and decreased symbols in blue.

## Data Availability

All data are contained within this manuscript or included in the supplemental data. Patient information was deidentified before analysis.

## Study Approval

The study was approved by UCSF’s Institutional Review Board and all research was performed in accordance with relevant guidelines/regulations in accordance with the Declaration of Helsinki. All participants signed an informed consent form approved by the IRB. Only deidentified data was used for analysis.

## Author Contribution

AB, TCN, and SSI contributed to the design of the study. AB wrote the first draft of the manuscript with input from SSI, JDK. AB, JDK, TCN, OF and SSI revised the final manuscript. SSI collected patient data, OF provided and curated metabolomic data. AB and JDK extracted and analyzed the data. All authors read and approved the final manuscript.

## Acknowledgments

The authors thank Drs. Anne Richards, Maddhu Rao, Aoife O’Donovan, and Lisa Talbot for sharing data sets and Thomas Metzler, and Erin Madden for help with clinical measures data. We acknowledge Maryann Lenoci and Leslie Ruoff, PGST for their logistical and technical support. This research was supported by Discovery Awards from the U.S. Army Medical Research & Materiel Command (USAMRMC), the Telemedicine & Advanced Technology Research Center (TATRC), at Fort Detrick, MD (SI: W81XWH-05-2-0094, W81XWH-15-PRMRP-DA, and SI: W81XWH-15-PRMRP-DA), and grants from the National Institute for Mental Health (TCN: 5R01MH073978-04, 5R34MH077667-03). Support was also provided by the Veterans Health Research Institute, the Mental Illness Research and Education Clinical Center of the US Veterans Health Administration, and the Clinical Research Center of the National Center for Advancing Translational Sciences, National Institutes of Health, through UCSF-CTSI Grant Number UL1 RR024131 and donor funds to AB.

